# High-level language brain regions are sensitive to sub-lexical regularities

**DOI:** 10.1101/2021.06.11.447786

**Authors:** Tamar I. Regev, Josef Affourtit, Xuanyi Chen, Abigail E. Schipper, Leon Bergen, Kyle Mahowald, Evelina Fedorenko

**Author notes:** **Corresponding Authors** Tamar I. Regev and Evelina Fedorenko, and; 43 Vassar Street, Room 46-3037, Cambridge, MA, 02139. Co-senior authors (Kyle Mahowald, Evelina Fedorenko). **Author contributions:** Conceptualization: TIR, LB, KM, EF. Design and materials creation and norming: AES, LB, KM, EF. Experimental script creation: KM. fMRI data collection: JA, KM, EF. fMRI data preprocessing and analysis: TIR, JA, XC, EF. Formal statistical analysis: TIR. Writing: TIR, KM, EF (the rest of the authors provided edits and comments). Overall project supervision: EF.

## Abstract

A network of left frontal and temporal brain regions supports ‘high-level’ language processing— including the processing of word meanings, as well as word-combinatorial processing—across presentation modalities. This ‘core’ language network has been argued to store our knowledge of words and constructions as well as constraints on how those combine to form sentences. However, our linguistic knowledge additionally includes information about sounds (phonemes) and how they combine to form clusters, syllables, and words. Is this knowledge of phoneme combinatorics also represented in these language regions? Across five fMRI experiments, we investigated the sensitivity of high-level language processing brain regions to sub-lexical linguistic sound patterns by examining responses to diverse nonwords—sequences of sounds/letters that do not constitute real words (e.g., punes, silory, flope). We establish robust responses in the language network to visually (Experiment 1a, n=605) and auditorily (Experiments 1b, n=12, and 1c, n=13) presented nonwords relative to baseline. In Experiment 2 (n=16), we find stronger responses to nonwords that obey the phoneme-combinatorial constraints of English. Finally, in Experiment 3 (n=14) and a post-hoc analysis of Experiment 2, we provide suggestive evidence that the responses in Experiments 1 and 2 are not due to the activation of real words that share some phonology with the nonwords. The results suggest that knowledge of phoneme combinatorics and representations of sub-lexical linguistic sound patterns are stored within the same fronto-temporal network that stores higher-level linguistic knowledge and supports word and sentence comprehension.

## INTRODUCTION

Languages contain rich statistical patterns across a range of information scales: from inter-word dependencies, to meanings of individual words/morphemes, to patterns of sounds within words. Traditionally, syntactic/combinatorial, lexical-semantic, and phonological representations and processes were construed as separate and assumed to be processed by distinct mechanisms (Chomsky, 1965, 1995; e.g., Chomsky and Halle, 1965; Pinker, 1991). However, in recent decades, linguistic theorizing has evolved toward a more integrated view of the language system, without sharp boundaries between our knowledge and processing of the sentence structure, word meanings, and sub-lexical sound patterns (Gaskell and Marslen-Wilson, 1997; Bybee, 1999, 2013; Jackendoff, 2007; Huettig et al., 2020; Jackendoff and Audring, 2020).

Evidence for links between sound patterns and the lexicon comes from corpus investigations across diverse languages that have observed that more frequent words are more *phonotactically regular*, i.e., obey the phoneme-combinatorial constraints of the language (e.g., Zipf, 1936; Landauer and Streeter, 1973; Frauenfelder et al., 1993; Mahowald et al., 2018; Pimentel et al., 2020), phonological clustering may be one organizing principle of the lexicon (e.g., Dautriche et al., 2017), and some sounds appear to be associated with aspects of meaning (e.g., Iwasaki et al., 2007; Monaghan et al., 2014; Larsson, 2015; Blasi et al., 2016; Winter et al., 2017; Sidhu and Pexman, 2018; Pimentel et al., 2019; Vinson et al., 2021). These links may be particularly important for language acquisition as every word we acquire is initially just a sequence of sounds (e.g., Davis et al., 2009; Perry et al., 2018; Jones et al., 2021).

Does this mean that—at the implementation level—the system that processes words and sentences also processes sub-lexical sound patterns? Past neuroscience research has not provided a clear answer. Prior neuroimaging investigations have reported effects for phonological manipulations in diverse left-hemisphere brain areas, including superior temporal gyrus (e.g., Paulesu et al., 1993; Price et al., 1997; Graves et al., 2007, 2008; DeWitt & Rauschecker, 2012; Lopopolo et al., 2017; Scott & Perrachione, 2019), supramarginal gyrus (e.g., Paulesu et al., 1993; Celsis et al., 1999; Church et al., 2011; Weiss et al., 2018; Yen et al., 2019), and inferior frontal cortex (e.g., Paulesu et al., 1993; Demonet et al., 1994; Poldrack et al., 1999; Burton, 2001; Myers et al., 2009; Xie & Myers, 2018). Similarly, lesions in these different brain areas (e.g., Geva et al., 2011; Pillay et al., 2014), as well as their interruption by electric/magnetic stimulation (e.g., Devlin et al., 2003; Boatman, 2004; Hartwigsen et al., 2016) have been shown to lead to impairments on phonological tasks, like rhyme judgments, nonword repetition, or phoneme identification.

Some of the brain areas implicated in phonological processing appear to overlap with the ‘core’ language network—a set of left-lateralized frontal and temporal areas that selectively respond to linguistic input, visual or auditory (e.g., Fedorenko et al., 2011; Monti et al., 2012) and support the processing of word meanings and combinatorial semantic and syntactic processes (e.g., Fedorenko et al., 2010, 2020; Bautista and Wilson, 2016). However, inferences about shared vs. distinct neural mechanisms based on the similarity of gross anatomical locations across studies are problematic (e.g., Poldrack, 2006; Fedorenko, 2021). Furthermore, most past studies have employed tasks that require computations beyond those engaged during naturalistic language processing and may recruit domain-general executive resources (e.g., Diachek et al., 2020).

To provide a clearer answer about whether the system that supports lexical and word-combinatorial processing is sensitive to sub-lexical sound patterns, we functionally defined the language network, and then examined these regions’ responses to nonwords during relatively naturalistic reading/listening across five fMRI experiments. Nonwords—visual or auditory— elicited robust responses in the language network despite their lack of meaning and ability to combine into larger units like phrases. Further, more word-like nonwords elicited stronger responses, and we provide suggestive evidence that this effect is not due to the activation of lexical representations of real-word ‘neighbors’ suggesting that sub-lexical units are represented within the system that extracts meaning from linguistic input.

## METHODS

### Participants

#### Experiments 1, 2, and 3

In total, 620 individuals (age 18-71 mean 24.9 +-7.3; 358 (57.7%) females) from the Cambridge/Boston, MA community participated for payment across 5 fMRI experiments (n=605 in Experiment 1a, n=12 in Experiment 1b, n=13 in Experiment 1c, n=16 in Experiment 2, n=14 in Experiment 3, for a total of 660 scanning sessions; **Table 1**). 43 participants overlapped between Experiment 1a and other experiments (12, 14 and 14 between Experiment 1a and Experiments 1c, 2 and 3, respectively; **Table 1**) and 4 participants overlapped between Experiments 2 and 3. 558 participants (∼90%, see **Table 1** for numbers per experiment) were right-handed, as determined by the Edinburgh handedness inventory (Oldfield, 1971), or self-report; the remaining participants were either left-handed (n=40), ambidextrous (n=14), or missing handedness information (n=8; see Willems et al., 2014, for arguments for including left-handers in cognitive neuroscience experiments). All participants were native English speakers, and all gave informed consent to participate in our experiments in accordance with the requirements of MIT’s Committee on the Use of Humans as Experimental Subjects (COUHES).

**Table 1.** Details of participants in all experiments.

| Experiment | 1a | 1b | 1c | 2 | 3 |
| --- | --- | --- | --- | --- | --- |
| Participants | 605 | 12 | 13 | 16 | 14 |
| Females | 349 | 7 | 6 | 11 | 11 |
| Age (mean, std) (years) | 24.9 (7.3) | 23.2 (4) | 24.7 (6.7) | 22.2 (7.1) | 25.1 (7.6) |
| Left handed (ambidextrous) | 36 (13) | 0 (0) | 1 (0) | 2 (0) | 1 (2) |
| Participants overlapping with Experiment: |  |  |  |  |  |
| 1a |  | 0 | 12 | 14 | 14 |
| 1b |  |  | 0 | 0 | 0 |
| 1c |  |  |  | 0 | 0 |
| 2 |  |  |  |  | 4 |
| 3 |  |  |  |  |  |

### Design, materials, and procedure

#### All experiments – overview

In all experiments, we examined responses to nonwords in the high-level language system. Therefore, in all experiments each participant completed a visual language localizer task (see Fedorenko et al., 2010, Scott et al., 2017, Chen, Affourtit et al., 2021 for evidence that this localizer is robust to modality) that served to functionally identify the language system within each individual participant. Each scanning session further included one or more tasks, including the critical experimental task and, in most cases, other tasks for unrelated studies, for a total duration of approximately two hours. The purpose of Experiments 1a, 1b and 1c was to examine the general robustness of responses to nonwords—phonotactically well-formed but meaningless and unconnected sound/letter strings—within the language system, across the visual and auditory modalities. In Experiments 2 and 3, we examined how finer phonotactic characteristics of nonwords affected these responses. The critical tasks included: Experiment 1a: passive reading of lists of nonwords from the visual language localizer (Fedorenko et al., 2010); Experiment 1b: listening to lists of nonwords (intermixed with some function words) and deciding whether a probe nonword appeared in the trial; Experiment 1c: passive listening to lists of nonwords; Experiment 2: reading lists of nonwords that parametrically vary in their word-likeness and deciding whether a probe nonword appeared in the trial; and Experiment 3: reading lists of nonwords with low or high phonological neighborhood with an accompanying 1-back memory task (see below for details of all tasks).

#### Language network localizer

This task was originally designed to elicit robust responses in the high-level language system, as described in detail in Fedorenko et al. (2010) and subsequent studies from the Fedorenko lab (and is available for download from https://evlab.mit.edu/funcloc/). In this task, participants passively read sentences and lists of unconnected, pronounceable nonwords in a blocked design.

In one version of the localizer (used in Experiments 1a, 1c, 2, and 3), the sentences were constructed to vary in content and syntactic structure, and the nonwords were created using the Wuggy software (Keuleers & Brysbaert, 2010), to match their phonotactic properties to those of the words used in the *sentence* condition (all the materials for this and other experiments are available at https://osf.io/6c2y7/). A trial consisted of 12 words/nonwords (presented one at a time, for 450 ms each), preceded by a 100 ms pretrial fixation and followed by a line drawing of a hand pressing a button appearing for 400 ms and finally a blank screen appearing for 100 ms, for a total trial duration of 6 s. Participants were instructed to press a button when the hand icon appeared. The task was included in order to help participants remain alert. Each block consisted of 3 trials and lasted 18 s. Each run consisted of 16 experimental blocks (8 per condition) and 5 fixation blocks (14 s each), for a total duration of 358 s (5 min 58 s). Each participant performed two runs, with condition order counterbalanced across runs.

In another version of the localizer (used in Experiment 1b), the *sentences* and *nonwords* conditions were presented along with two other conditions (*Jabberwocky* sentences and lists of unconnected words) that are not relevant to the current study (Experiment 2 in Fedorenko et al. 2010). As in the version described above, the sentences were constructed to vary in content and syntactic structure. The *nonwords* condition contained nonwords and function words (the function words were included to match this condition to the *Jabberwocky* sentence condition, which uses function words to create English-like syntactic patterns). The nonwords were created by a) syllabifying all content words in the *sentence* condition, b) replacing single phonemes in monosyllabic words in a way that respects phonotactic constraints of English, c) recombining the syllables in a way so as to create words whose lengths are similar in distribution to those of the content words in the *sentence* condition, and, finally, d) assembling the resulting nonwords and the function words into 8-word/nonword-long strings while minimizing the possibility of any local syntactic structure building. A trial consisted of 8 words/nonwords (presented one at a time, for 350 ms each), followed by a 300 ms fixation, a memory probe appearing on the screen for 350 ms, a period of 1,000 ms during which participants were instructed to press one of two buttons to indicate whether the probe word/nonword appeared in the preceding trial, and a 350 ms fixation, for a total trial duration of 4.8 s. Only nonwords were used as probes in the *nonwords* condition and the correct response was YES in half of the trials. Each experimental block consisted of 5 trials and lasted 24 s. Each run consisted of 16 experimental blocks (4 per condition) and 5 fixation blocks (16 s each), for a total duration of 464 s (7 min 44 s). Each participant performed between 6-8 runs, with condition order counterbalanced across runs.

The *Sentences>Nonwords* contrast targets brain regions that support high-level language comprehension, including lexico-semantic and combinatorial processes (e.g., Fedorenko et al., 2010, 2020; Fedorenko, Nieto-Castañon, et al. 2012; Blank et al., 2016) and has been shown to be robust to changes in materials, task, timing parameters, and other aspects of the procedure (Fedorenko et al. 2010; Fedorenko 2014; Mahowald and Fedorenko 2016; Scott et al. 2017; Diachek, Blank, Siegelman et al., 2020).

#### Experiment 1a

##### Critical task – Passive reading of lists of nonwords from the language localizer

To examine the robustness of responses to visually presented nonwords in the language regions, we used the *nonwords* condition from the language localizer. Response magnitudes were estimated with cross-validation across experimental runs to ensure that the data used for the localization of the language regions was independent from the data used to estimate the responses to nonwords in this critical task (we first used run 1 to define the regions of interest and run 2 to estimate the responses; then used run 2 to define the regions and run 1 to estimate the responses; finally, we averaged the estimates across the two runs to derive a single estimate per participate per region; e.g., Kriegeskorte et al., 2011). As noted above, the nonwords were accompanied by a simple button-press task to maintain alertness (behavioral responses for this and all other experiments are summarized in a supplementary table available at https://osf.io/6c2y7/).

#### Experiment 1b

##### Critical task – Listening to lists of nonwords with a memory probe task

To examine the robustness of responses to auditorily presented nonwords in the language regions, we used the *nonwords* condition from an auditory language experiment that was published previously (Experiment 3 in Fedorenko et al., 2010). Participants listened to recordings of lists of nonwords (and materials from three other conditions not relevant to the current study) in a blocked design. The nonword lists were constructed by re-combining the syllables that comprised the words in the real-word conditions of the experiment (to preserve the phonotactic well-formedness) and recorded by a female native English speaker (see Fedorenko et al., 2013 for a detailed acoustic analysis of these materials). Example items are shown in **Figure 1**.

**Figure 1.**
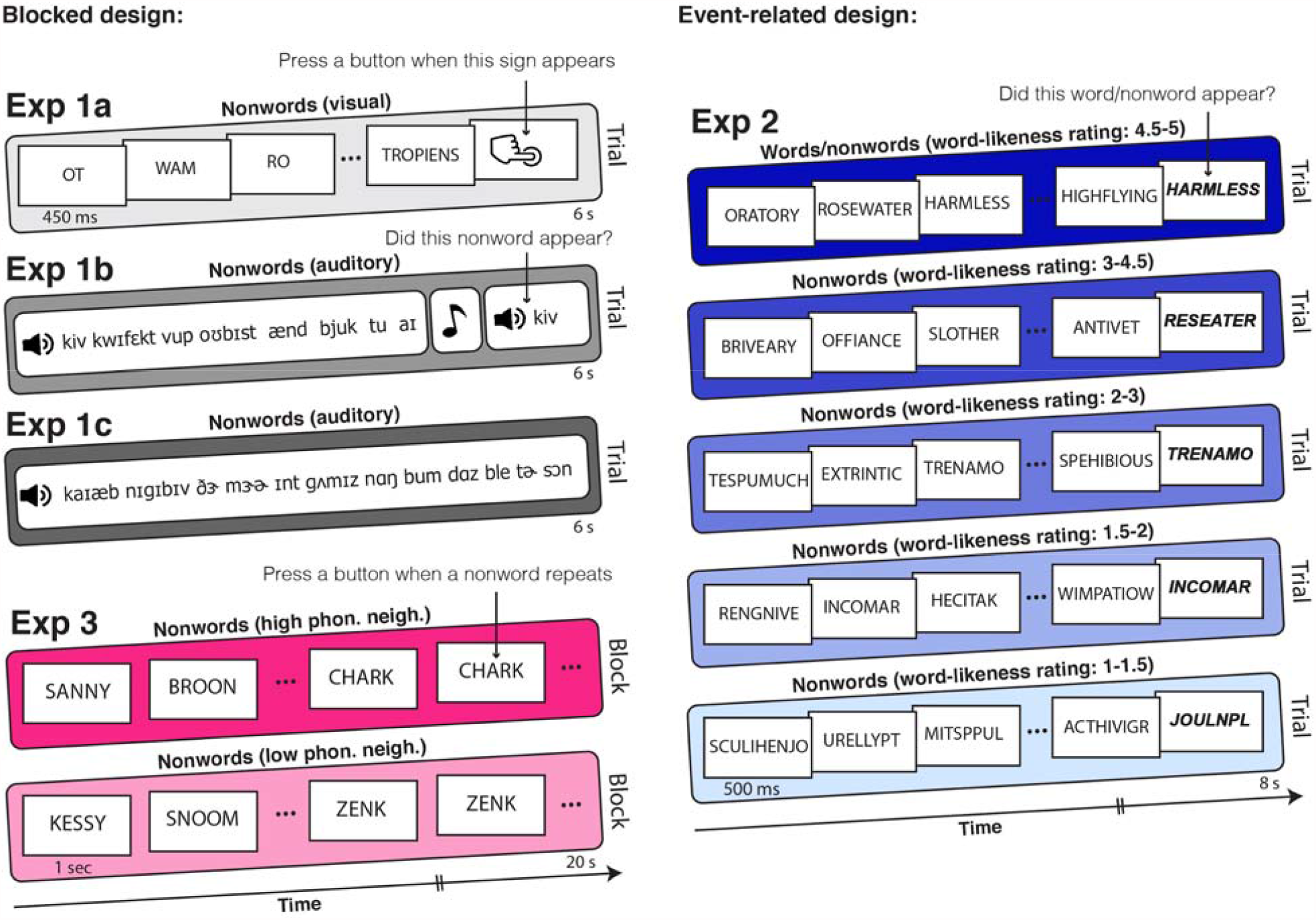
Procedure and example stimuli for all experiments. Color-filled rounded rectangles represent a typical block or trial in a specific condition of each experiment. The color codes match those used in Figure 2. Experiment numbers and conditions are indicated above each rectangle. Left column – experiments with a blocked design, right column – experiments with an event-related design. **Left to right, top to bottom** (see methods for further detail): **Exp 1a –** Passive reading of lists of nonwords from the language localizer. **Exp 1b –** Listening to lists of nonwords with a memory probe task. **Exp 1c** – Passive listening to lists of nonwords. **Exp 3 –** Reading of lists of nonwords with a low or high phonological neighborhood, with a 1-back task. **Exp 2 –** Reading of lists of nonwords parametrically varying in word-likeness, with a memory probe task.

A trial consisted of a recording of 8 nonwords (the strings consisted of 3-7 pronounceable nonwords and 1-5 function words, such as ‘a’, ‘the’, ‘of’, ‘with’, which were included in order to match this condition to another condition in the original study; the order of nonwords and function words did not allow for any syntactic structure building) that lasted 3300-4300 ms, followed by a beep tone (100 ms) and then an auditory memory probe (presented for up to 1000 ms). Participants were instructed to indicate whether or not the probe appeared in the preceding list by pressing 1 of 2 buttons during the period following the memory probe, which lasted until the end of the trial, for a total trial duration of 6 s. The memory probes came from the preceding stimulus on half of the trials, and were approximately uniformly distributed across the 8 positions. Incorrect probes were the shuffled correct probes from other lists.

Each block consisted of 4 trials and lasted 24 s. Each run consisted of 16 experimental blocks (4 per condition) and 5 fixation blocks (16 s each), for a total duration of 464 s (7 min 44 s). All but one participant completed 4 runs for a total of 16 blocks per condition; the remaining participant completed 7 runs for a total of 28 blocks per condition. Condition order was palindromic within a run and counterbalanced across runs.

#### Experiment 1c

##### Critical task – Passive listening to lists of nonwords

To replicate and generalize the results from Experiment 1b, we used a *nonwords* condition from another auditory experiment (Experiment 4 in Chen, Affourtit et al., 2021). Participants passively listened to recordings of lists of nonwords (and materials from several other conditions not relevant to the current study) in a blocked design. The nonword lists were constructed by taking a set of sentences and replacing each word with a nonword that has a similar phonological structure but that does not have any meaning and recorded by a female and a male native English speakers (in the experiment, half of the trials came from the female speaker, and the other half from the male speaker). Example items are shown in **Figure 1**.

A trial consisted of a recording of 12 nonwords that lasted 5-5.95 s followed by a brief 0.05s silence. Recordings that were shorter than 5.95s were padded with silence, and recordings that were longer than 5.95 seconds were trimmed off at 5.95s. Participants were instructed to listen attentively.

Each block consisted of 3 trials and lasted 18 s. Each run consisted of 18 experimental blocks (2 per condition) and 3 fixation blocks (14 s each), for a total duration of 366s (6min 6s). Condition order was palindromic within a run and counterbalanced across runs. Most participants (n=9) completed 6 runs (for a total of 12 blocks per condition), one subject completed 5 runs (for a total of 10 blocks per condition), one subject completed 7 runs (for a total of 14 blocks per condition), and the remaining two subjects completed 8 runs (for a total of 16 blocks per condition).

#### Experiment 2

##### Critical task – Reading of lists of nonwords parametrically varying in word-likeness, with a memory probe task

This experiment was designed to probe the responses in the language system to words and nonwords that vary in their degree of word-likeness, to test whether more word-like nonwords elicit a stronger response. Participants read lists of words and nonwords (and materials from five other conditions not relevant to the current study) in an event-related design. The nonwords were created from real words via one or multiple letter replacements, as detailed below. Original words and different resulting versions of nonwords were grouped into 5 conditions based on the magnitude of word-likeness ratings obtained in a behavioral norming study (conducted online, with different participants). Example items are shown in **Figure 1**.

A trial consisted of 12 words/nonwords (presented one at a time, for 500 ms each), followed by a blank screen presented for 300 ms, followed by a memory probe presented in blue font for 1200 ms, followed again by a blank screen for 500 ms for a total trial duration of 8 s. Participants were instructed to indicate whether or not the probe appeared in the preceding list by pressing 1 of 2 buttons from the moment of probe appearance. The memory probes came from the preceding stimulus on half of the trials, and were approximately uniformly distributed across the 12 positions. Incorrect probes were the shuffled correct probes from other sequences in the same condition.

Each run consisted of 50 trials (5 per condition) and 80 s of fixation, for a total duration of 480 s (8 min). The optseq2 algorithm (Dale, 1999) was used to create condition orderings and to distribute fixation among the trials so as to optimize our ability to deconvolve responses to the different conditions. Condition order varied across runs and participants. Most participants (n=13) completed 5 runs (for a total of 25 trials per condition); the remaining 3 completed 4 or 3 runs (for a total of 20 or 15 trials, respectively) due to time constraints.

#### Construction and norming of the materials

To create the nonwords, a large set of real trisyllabic words (n=20695) was first identified. For each word, 14 versions of nonwords were created by replacing random letters with other letters, while ensuring that the local trigram context (the letter preceding the critical letter, the critical replaced letter, and the letter following it) is attested in English. For example, consider the word “BLACKBERRY”; the letter C could be replaced with the letter R because the string “ARK” is attested (e.g., BARK), or with the letter L because the string “ALK” is attested (e.g., ALKALINE), but not with the letter X because the string “AXK” is not attested. This replacement process was repeated on the resulting nonword (e.g., BLARKBERRY in this example) using the same constraints, up to 14 times total. This procedure resulted in a set of 310,425 words and nonwords including the original words and all the resulting nonwords from the 14 letter-replacement iterations done on each word. A subset of these materials (n=900, sampled ∼equally from the 15 ‘levels’, i.e., number of replaced letters, between 0 and 14) were presented to participants via Amazon.com’s Mechanical Turk platform. Participants were presented with one word/nonword at a time and asked to rate each for how “word-like” it was, on a scale from 1 (not at all word-like) to 5 (very word-like). The words/nonwords were then divided into 5 groups according to the word-likeness ratings, which were accordingly binned as follows, from least to most word-like: 1-1.5, 1.5-2, 2-3, 3-4.5, 4.5-5. The binning was determined to balance the number of items within each group (condition). Each group consisted of 180 items, except for the most word-like group, for which there were only 173 items. The most word-like group consisted of mostly real words and all the other 4 groups consisted of only nonwords. 12-item strings were created from these materials for each condition.

#### Experiment 3

##### Critical task – Reading of lists of nonwords with a low or high phonological neighborhood, with a 1-back task

This experiment was designed to probe the responses in the language system to nonwords varying in their phonological neighborhood, to test whether nonwords with a high phonological neighborhood elicit a stronger response (due to activating real word neighbors). Participants read lists of nonwords in a blocked design. The nonwords were created as detailed below.

A block consisted of 20 nonwords (presented one at a time, for 1 s each) for a total block duration of 20s. Participants were instructed to read the nonwords and to press a button when a nonword is repeated in succession (1-back task). Nonword repetitions occurred 4 times per block, and were approximately uniformly distributed across the block. Each run consisted of 10 experimental blocks (5 per condition) and 3 fixation blocks (14 s each), for a total duration of 242 s (4 min 2 s). Condition order varied across runs and participants. Most participants (n=10) performed 3 runs (for a total of 15 blocks per condition); the remaining 4 participants performed 2 runs (for a total of 10 blocks per condition) due to time constraints.

#### Construction of the materials

To create the nonwords, a 3-gram model over phonemes was used, using the generative procedure described in Dautriche, et al. (2017). The logic of this n-gram model of phonotactics is that each phoneme is generated probabilistically, conditioned on the preceding two phonemes.

Using this model, a large candidate set of candidate nonwords was sampled without replacement. Then, 80 pairs of nonwords were selected such that they were matched on consonant-vowel patterns (i.e. length in letters and number of syllables), and the two sets of nonwords were roughly matched on a pronunciation-based phonotactic (‘BLICK’) score (Hayes & Wilson 2008; Hayes 2012). A 2-sample t-test of BLICK scores between the groups showed no significant difference (t(158)=0.05, p=0.96). But the groups were constructed to maximally differ with respect to phonological neighborhood size (the number of real English words that are 1 edit away from the nonword; for example, phonological neighbors of the nonword “ZAT” include “BAT”, “CAT”, and “ZAP”, among others). In the high neighborhood group, each nonword had at least 9 neighbors (mean=11, SD=2.6), and in the low neighborhood group, each nonword had at most 3 neighbors (mean=1.85, SD=0.9, 2-sample t-test of neighborhood scores between the groups: t(158)=28.7, p<0.0001).

### Phonotactic probability and orthographic neighborhood size of stimuli in Experiments 2 and 3

To investigate which features of the nonword stimuli may contribute to neural responses in the language system, we further calculated the phonotactic probability and orthographic neighborhood size of the (visually presented) stimuli in Experiments 2 and 3. We used the English Lexicon Project website https://elexicon.wustl.edu/, Balota et al., 2007) for both measures. This website allows one to submit lists of written nonwords and outputs a series of characteristics calculated based on an English corpus (Balota et al., 2007). For the phonotactic probability measure, we used the mean bigram frequency, which is the sum of bigram counts (where a bigram, here, is a sequence of two letters like CA and AT in CAT) of all the local bigrams within a nonword, divided by the number of bigrams. The orthographic neighborhood size measure corresponds to the number of real words that can be obtained by changing one letter while preserving the identity and positions of the other letters (i.e., Coltheart’s N; Coltheart, Davelaar, Jonasson, & Besner, 1977). We chose to use an orthographic neighborhood measure and not a phonological one (as we originally did for designing the materials for Experiment 3) because the pronunciation of many nonwords is inherently ambiguous (e.g., would the nonword

KLOUGH be pronounced to rhyme with *through, trough*, or *tough*?). In general, though, we believe that our results are robust to whether we use orthographic or phonological neighbors, which we verify in Experiment 3 (R=0.55, relative to no correlation p=2.6E-14). Having obtained the phonotactic probability and orthographic neighborhood measures for each nonword presented in Experiments 2 and 3, we then calculated the average and standard error across all nonwords in each condition (five conditions varying in word-likeness in Experiment 2, and two conditions varying in phonological neighborhood in Experiment 3).

### fMRI data acquisition, preprocessing, and first-level modeling

#### Experiments 1a, 1c, 2 and 3

##### Data acquisition

Whole-brain structural and functional data were collected on a whole-body 3 Tesla Siemens Trio scanner with a 32-channel head coil at the Athinoula A. Martinos Imaging Center at the McGovern Institute for Brain Research at MIT. T1-weighted structural images were collected in 176 axial slices with 1 mm isotropic voxels (repetition time (TR)=2530 ms; echo time (TE)=3.48 ms). Functional, blood oxygenation level-dependent (BOLD) data were acquired using an EPI sequence with a 90° flip angle and using GRAPPA with an acceleration factor of 2; the following parameters were used: thirty-one 4.4 mm thick near-axial slices acquired in an interleaved order (with 10% distance factor), with an in-plane resolution of 2.1 mm × 2.1 mm, FoV in the phase encoding anterior to posterior (A>>P) direction 200 mm and matrix size 96 × 96 voxels, TR=2,000 ms and TE=30 ms. The first 10 s of each run were excluded to allow for steady state magnetization.

#### Experiment 1b

##### Data acquisition

Structural and functional data were collected on the whole-body 3 Tesla Siemens Trio scanner at the Athinoula A. Martinos Imaging Center at the McGovern Institute for Brain Research at MIT. T1-weighted structural images were collected in 128 axial slices with 1.33 mm isotropic voxels (TR=2000 ms, TE=3.39 ms). Functional, blood-oxygenation-level-dependent (BOLD) data were acquired in 3.1 × 3.1 × 4 mm voxels (TR=2000 ms, TE=30 ms) in 32 near-axial slices. The first4sof each run were excluded to allow for steady state magnetization.

#### All Experiments

##### Preprocessing

Data preprocessing was carried out with SPM12 (using default parameters, unless specified otherwise) and supporting, custom MATLAB scripts. Preprocessing of functional data included motion correction (realignment to the mean image using 2^nd^-degree b-spline interpolation), normalization into a common space (Montreal Neurological Institute (MNI) template) (estimated for the mean image using trilinear interpolation), resampling into 2 mm isotropic voxels, smoothing with a 4 mm FWHM Gaussian filter and high-pass filtering at 128 s.

##### First-level modeling

For both the language localizer task and the critical experiments, a standard mass univariate analysis was performed in SPM12 whereby a general linear model (GLM) estimated, for each voxel, the effect size of each condition in each experimental run. These effects were each modeled with a boxcar function (representing entire blocks/events) convolved with the canonical Hemodynamic Response Function (HRF). The model also included first-order temporal derivatives of these effects, as well as nuisance regressors representing entire experimental runs and offline-estimated motion parameters.

### Definition of the language functional regions of interest (fROIs) (all Experiments)

For each critical task, we defined first a set of language functional ROIs using group-constrained, subject-specific localization (Fedorenko et al 2010). In particular, each individual map for the *sentences>nonwords* contrast from the language localizer was intersected with a set of five binary masks. These masks (**Figure 1**; available at http://web.mit.edu/evlab//funcloc/#parcels and OSF https://osf.io/6c2y7/ were derived from a probabilistic activation overlap map for the same contrast in a large set of participants (n=220) using watershed parcellation, as described in

Fedorenko et al. (2010) for a smaller set of participants. These masks covered the fronto-temporal language network in the left hemisphere (we excluded the AngG parcel because the AngG fROI has been shown to consistently pattern differently from the rest of the language network across diverse measures (e.g., Blank et al., 2014; Chai et al., 2016; Mineroff, Blank et al., 2018; Pritchett et al., 2018), including responding more strongly to visual meaningful stimuli than to sentences (Amit et al., 2017; Ivanova et al., 2021), which suggests that it is not a language region). Within each mask, a participant-specific language fROI was defined as the top 10% of voxels with the highest *t*-values for the localizer contrast.

## Statistical analyses

### Validation of the language fROIs (all Experiments)

To ensure that the language fROIs behave as expected (i.e., show a reliably greater response to the sentences condition compared to the *nonwords* condition), we used an across-runs cross-validation procedure (e.g., Nieto-Castañón and Fedorenko, 2012). In this analysis, the first run of the localizer was used to define the fROIs, and the second run to estimate the responses (in percent BOLD signal change, PSC, relative to fixation baseline) to the localizer conditions, ensuring independence (e.g., Kriegeskorte et al., 2009); then the second run was used to define the fROIs, and the first run to estimate the responses; finally, the extracted magnitudes were averaged across the two runs to derive a single response magnitude for each of the localizer conditions. Statistical analyses were performed on these extracted PSC values. Namely, for each of the five language fROIs identified, we fit a linear mixed-effect regression model, predicting the level of PSC to sentences relative to nonwords. The model included fixed effects for an intercept and a slope variable encoding the difference between sentences and nonwords on top of the common intercept. This scheme was implemented by coding sentences as a +0.5 factor and nonwords as a −0.5 factor. The model additionally included random terms for both the intercept and the slope variable encoding the difference between sentence and nonwords, both grouped by participant:

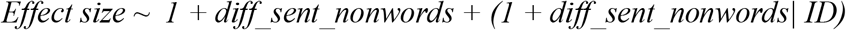

Where *1* denotes the intercept, *diff_sent_nonwords* denotes the difference between *sentence* and *nonwords* slope variable, encoded as explained above, and *ID* denotes a unique identification number per participant.

In this coding scheme, the intercept estimate reflects the average PSC response for the sentence and nonword conditions together and the slope variable estimate reflects the difference between the *sentence* and *nonwords* conditions. Therefore, to test the validity of the language fROIs, we examined the values of the fixed intercept and slope variable estimates. Both of these estimates had to be significantly positive. The results were FDR-corrected for the 5 ROIs.

### Analyses of the critical tasks (all experiments)

To estimate the responses in the language fROIs to the conditions of the critical tasks, in each experiment the data from all the runs of the language localizer were used to define the fROIs, and the responses to each condition were then estimated in these regions. The critical conditions were as follows (see *Design, materials, and procedure* above): i) in Experiment 1a: visual nonwords from the language localizer, ii) in Experiment 1b: auditory words/nonwords, iii) in Experiment 1c: auditory nonwords, iv) in Experiment 2: five word/nonword conditions parametrically varying in ‘word-likeness’, and v) in Experiment 3: two nonword conditions with low or high phonological neighborhood.

For each experiment, we used linear mixed-effect (LME) regression models (using Matlab *fitlme* routine) to determine the significance of activations of the critical conditions within the language network. We used these models in two ways: i) to examine the response within the language network as a whole; and ii) to examine the responses in each of the 5 language fROIs separately. Treating the language network as an integrated system is reasonable given that the regions of this network a) show similar functional profiles, both with respect to selectivity for language over non-linguistic processes (e.g., Fedorenko et al., 2011) and with respect to their role in lexico-semantic and syntactic processing (e.g., Blank et al., 2016; Fedorenko et al., 2020), and b) exhibit strong inter-region correlations in both their activity during naturalistic cognition paradigms (e.g., Blank et al., 2014; Braga et al., 2020) and key functional markers, like the strength of response or the extent of activation in response to language stimuli (e.g., Mahowald and Fedorenko, 2016; Mineroff, Blank et al., 2018; Affourtit, Lipkin et al. in prep.). However, because we wanted to allow for the possibility that language regions might differ in their response to nonwords, as well as in order to examine the robustness of the effects across the language fROIs, we supplement the network-wise analyses with the analyses of the five language fROIs separately.

For each of the five language fROIs identified, we fit a linear mixed-effect regression model, predicting the level of PSC in the target language fROI in the contrasted conditions.

In the case of modeling a condition with a single level, as in Exp1a, b and c, which all contained a single critical condition (nonwords), this condition was modeled as the intercept of the model. The intercept estimates are reported as representing the condition. The model then included a fixed effect for the intercept, and a random intercept grouped by participant.

For the network-level analysis, we included a random intercept grouped by ROI:

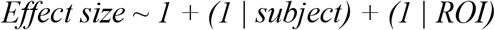

For the ROI-level analysis, we ran this model for each ROI:

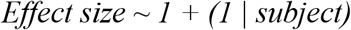

The p-values (comparing the intercept estimate to 0) were FDR-corrected for the 5 ROIs.

In the case of modeling a condition with multiple levels, we added a slope variable encoding the effect of the critical condition beyond the common intercept. For Exp 2, we modeled the 5 levels of word-likeness by coding them on a linear scale from −2 to 2 (multiplying brain activity by the factors −2, −1, 0, 1, 2) from low to high word-likeness, respectively. In Exp 3, we coded the low phonological neighborhood condition as −0.5 and the high neighborhood condition as 0.5.

In these cases, the model included fixed effects for the intercept and condition (the slope variable coding the critical condition) and potentially correlated random intercepts and slopes grouped by participant. Here, the intercept represents the mean brain activity across all the levels of the critical condition and the condition slope estimate represents the deviation in brain activity due to the different levels of the critical conditions. Therefore, the overall effect of the critical condition was significant if the condition estimates were significantly different from 0.

For the network-level analysis for Exp 2 and 3 we included potentially correlated random intercept and slopes grouped by fROI:

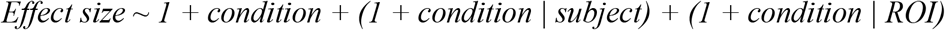

For the ROI-level analysis, we ran this model for each ROI:

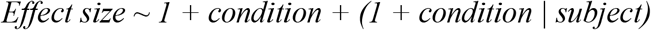

The p-values comparing the condition estimates to 0 were FDR-corrected for the 5 ROIs.

## RESULTS

### Validation of the language fROIs (all experiments)

In all experiments, each of the five language functional regions of interest (fROIs, see **Figure 2B** for parcel locations) showed a reliably above-baseline response to *sentences* (all intercept estimates>0, *p*s<0.001, FDR-corrected for the 5 fROIs; full results available at OSF: https://osf.io/6c2y7/), as well as a robust *sentences>nonwords* effect (all slope estimates>0, *p*s<0.001, FDR-corrected for the 5 fROIs), consistent with much previous work (e.g., Fedorenko et al., 2010; Mahowald and Fedorenko, 2016; Diachek, Blank, Siegelman et al., 2020).

**Figure 2.**
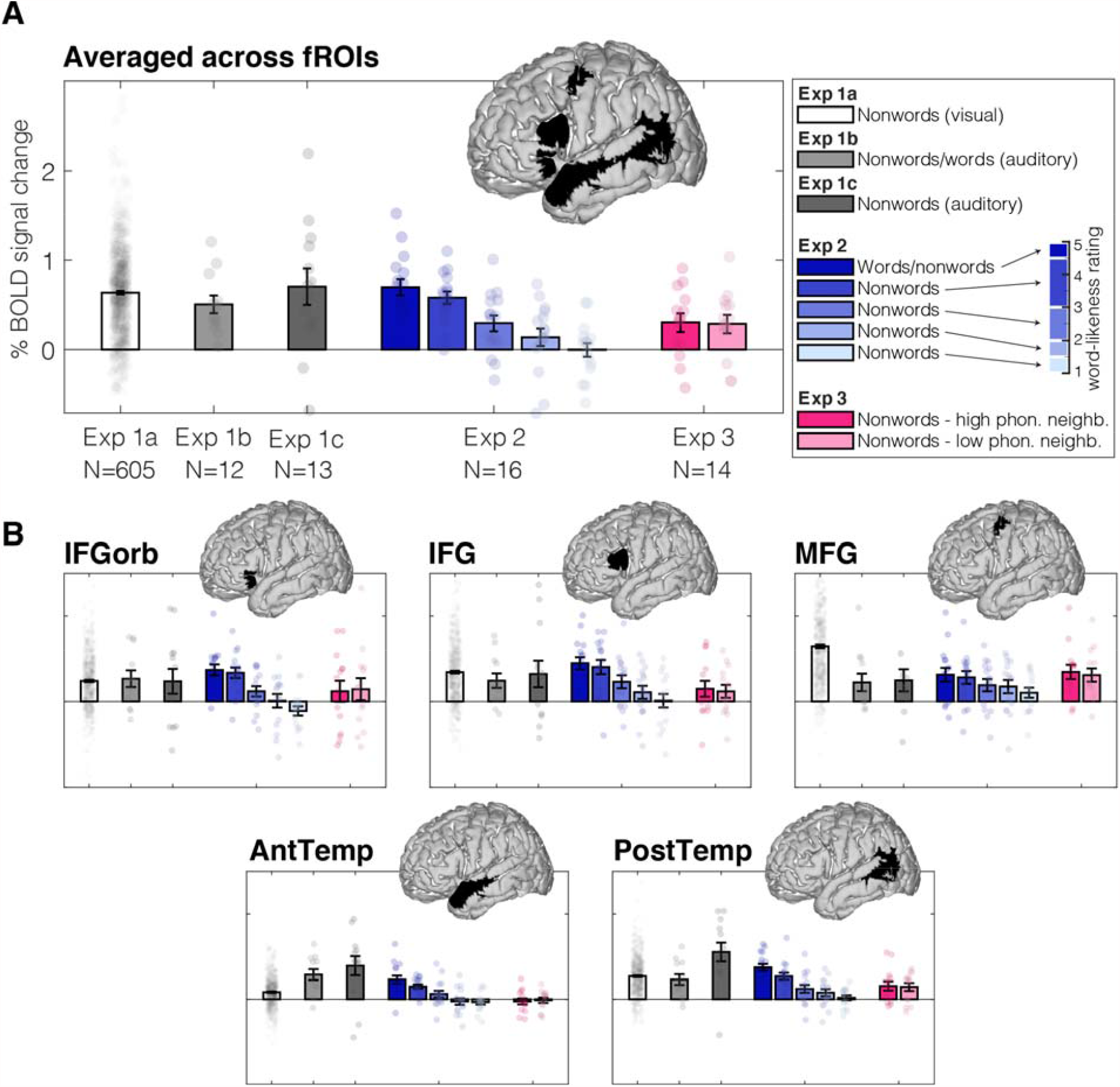
Sensitivity of the language system to nonwords in all experiments. Bargraphs are % BOLD signal change relative to a fixation baseline averaged across participants in each specific experiment (number of participants specified on abscissa). Dots are individual participants. The brain images display the parcels used to define the fROIs; individual fROIs are 10% of most language-responsive voxels within each parcel. **A** – Responses in all five language fROIs together. **B** – Responses in each fROI separately. IFGorb – inferior frontal gyrus orbital, IFG – inferior frontal gyrus, MFG – medial frontal gyrus, AntTemp – anterior temporal, PostTemp – posterior temporal.

### Key result 1: The language fROIs respond robustly to visually and auditorily presented nonwords (Experiments 1a-c)

In Experiment 1a, visually presented nonwords elicited a robust response relative to the fixation baseline across the language network, treating the fROIs as a random effect (*p*<0.001; **Table 3, Figure 2A**), as well as in each of the five language fROIs individually (*p*s<0.001, FDR-corrected; **Table 2, Figure 2B**). The behavioral responses revealed that participants remained alert during the task (87% responses, SD=27%, RT=427ms, SD=96ms). For this and other experiments, raw behavioral data are available at OSF: https://osf.io/6c2y7/.

**Table 2.** Linear mixed effects results for each fROI separately.

| Experiment | ROI | Fixed Effect | Estimate | SE | tStat | DF | pValue | p_FDR | H_FDR |
| --- | --- | --- | --- | --- | --- | --- | --- | --- | --- |
| 1a | IFGorb | Intercept | 0.48 | 0.03 | 18.49 | 604 | 8.3E-61 | 1.0E-60 | 1 |
| 1a | IFG | Intercept | 0.69 | 0.03 | 23.62 | 604 | 7.3E-88 | 1.2E-87 | 1 |
| 1a | MFG | Intercept | 1.29 | 0.04 | 29.86 | 604 | 4.8E-121 | 2.4E-120 | 1 |
| 1a | AntTemp | Intercept | 0.17 | 0.01 | 11.26 | 604 | 7.6E-27 | 7.6E-27 | 1 |
| 1a | PostTemp | Intercept | 0.54 | 0.02 | 28.82 | 604 | 1.6E-115 | 3.9E-115 | 1 |
| 1b | IFGorb | Intercept | 0.54 | 0.19 | 2.88 | 11 | 1.5E-02 | 1.9E-02 | 1 |
| 1b | IFG | Intercept | 0.49 | 0.16 | 3.01 | 11 | 1.2E-02 | 2.0E-02 | 1 |
| 1b | MFG | Intercept | 0.45 | 0.20 | 2.30 | 11 | 4.2E-02 | 4.2E-02 | 1 |
| 1b | AntTemp | Intercept | 0.57 | 0.12 | 4.75 | 11 | 6.0E-04 | 3.0E-03 | 1 |
| 1b | PostTemp | Intercept | 0.46 | 0.12 | 3.81 | 11 | 2.9E-03 | 7.2E-03 | 1 |
| 1c | IFGorb | Intercept | 0.47 | 0.28 | 1.70 | 12 | 1.2E-01 | 1.2E-01 | 0 |
| 1c | IFG | Intercept | 0.65 | 0.30 | 2.17 | 12 | 5.1E-02 | 8.5E-02 | 0 |
| 1c | MFG | Intercept | 0.50 | 0.25 | 2.01 | 12 | 6.7E-02 | 8.4E-02 | 0 |
| 1c | AntTemp | Intercept | 0.78 | 0.21 | 3.65 | 12 | 3.4E-03 | 8.4E-03 | 1 |
| 1c | PostTemp | Intercept | 1.10 | 0.21 | 5.23 | 12 | 2.1E-04 | 1.1E-03 | 1 |
| 2 | IFGorb | Intercept | 0.29 | 0.09 | 3.30 | 78 | 1.5E-03 | 1.8E-03 | 1 |
| 2 | IFGorb | Slope | 0.26 | 0.04 | 6.16 | 78 | 3.0E-08 | 3.8E-08 | 1 |
| 2 | IFG | Intercept | 0.48 | 0.13 | 3.61 | 78 | 5.4E-04 | 1.3E-03 | 1 |
| 2 | IFG | Slope | 0.23 | 0.03 | 8.63 | 78 | 5.6E-13 | 1.4E-12 | 1 |
| 2 | MFG | Intercept | 0.43 | 0.13 | 3.30 | 78 | 1.5E-03 | 2.4E-03 | 1 |
| 2 | MFG | Slope | 0.11 | 0.03 | 4.21 | 78 | 6.9E-05 | 6.9E-05 | 1 |
| 2 | AntTemp | Intercept | 0.15 | 0.05 | 2.86 | 78 | 5.4E-03 | 5.4E-03 | 1 |
| 2 | AntTemp | Slope | 0.14 | 0.02 | 6.96 | 78 | 9.2E-10 | 1.5E-09 | 1 |
| 2 | PostTemp | Intercept | 0.34 | 0.06 | 5.47 | 78 | 5.3E-07 | 2.7E-06 | 1 |
| 2 | PostTemp | Slope | 0.18 | 0.02 | 10.86 | 78 | 2.8E-17 | 1.4E-16 | 1 |
| 3 | IFGorb | Intercept | 0.27 | 0.24 | 1.12 | 26 | 2.7E-01 | 3.4E-01 | 0 |
| 3 | IFGorb | Slope | -0.05 | 0.12 | -0.42 | 26 | 6.8E-01 | 1.1E+00 | 0 |
| 3 | IFG | Intercept | 0.27 | 0.16 | 1.69 | 26 | 1.0E-01 | 1.7E-01 | 0 |
| 3 | IFG | Slope | 0.06 | 0.09 | 0.68 | 26 | 5.0E-01 | 1.3E+00 | 0 |
| 3 | MFG | Intercept | 0.66 | 0.15 | 4.32 | 26 | 2.0E-04 | 1.0E-03 | 1 |
| 3 | MFG | Slope | 0.07 | 0.06 | 1.29 | 26 | 2.1E-01 | 1.0E+00 | 0 |
| 3 | AntTemp | Intercept | -0.04 | 0.05 | -0.64 | 26 | 5.2E-01 | 5.2E-01 | 0 |
| 3 | AntTemp | Slope | -0.03 | 0.07 | -0.40 | 26 | 6.9E-01 | 8.7E-01 | 0 |
| 3 | PostTemp | Intercept | 0.30 | 0.09 | 3.29 | 26 | 2.9E-03 | 7.2E-03 | 1 |
| 3 | PostTemp | Slope | 0.02 | 0.06 | 0.33 | 26 | 7.4E-01 | 7.4E-01 | 0 |

**Table 3.** Linear mixed-effects results for the whole language system together

| Experiment | FixedEffectName | Estimate | SE | tStat | DF | pValue | H |
| --- | --- | --- | --- | --- | --- | --- | --- |
| 1a | Intercept | 0.63 | 0.17 | 3.82 | 3024 | 1.4E-04 | 1 |
| 1b | Intercept | 0.50 | 0.09 | 5.29 | 59 | 1.9E-06 | 1 |
| 1c | Intercept | 0.70 | 0.21 | 3.33 | 64 | 1.5E-03 | 1 |
| 2 | Intercept | 0.34 | 0.08 | 4.02 | 398 | 7.0E-05 | 1 |
| 2 | Slope | 0.18 | 0.03 | 6.38 | 398 | 4.9E-10 | 1 |
| 3 | Intercept | 0.29 | 0.13 | 2.20 | 138 | 3.0E-02 | 1 |
| 3 | Slope | 0.02 | 0.09 | 0.18 | 138 | 8.6E-01 | 0 |

Similarly, auditorily presented nonwords elicited a robust response relative to the fixation baseline across the language network, treating the fROIs as a random effect, in both Experiment 1b and Experiment 1c (*p*s<0.01; **Table 3, Figure 2A**). In Experiment 1b, the effect was reliable in each of the five language fROIs (*p*s<0.05, FDR-corrected, **Table 2, Figure 2B**), and in Experiment 1c, the effect was reliable in the two temporal fROIs (AntTemp and PostTemp fROIs; *p*s<0.01, FDR-corrected; **Table 2, Figure 2B**), with the three frontal fROIs showing a trend (IFGorb, IFG, and MFG fROIs; 0.05<*p*s<0.09, FDR-corrected; **Table 2, Figure 2B**). The average performance in the memory task was 82.2% correct responses (SD=1.4%).

Thus, Experiments 1a-1c revealed robust sensitivity in the language fROIs to nonwords across modalities and tasks.

### Key result 2: The language fROIs respond more strongly to more word-like nonwords (Experiment 2)

The word-likeness manipulation resulted in a gradient of fMRI response strength in the language network such that more word-like nonwords/words elicited stronger responses (*p*<0.001, **Table 2, Figure 2**). This effect was reliable in each of the five language fROIs (*ps*<0.001, FDR-corrected, **Table 3**), and as well when considering the language system together, treating the fROIs as a random grouping factor (all *ps*<0.001, FDR-corrected, **Table 2**).

Behavioral performance on the memory probe task was relatively high across conditions (70-80% on average across the word-likeness conditions, **Figure 3C**).Word-likeness had a small effect on performance, with better performance for more word-like conditions (fixed slope of accuracy as a function of word-likeness in a LME model including fixed and random intercept and slope grouped by participant; −0.02, t(78)= −2.23, p=0.025, **Figure 3C**).

**Figure 3.**
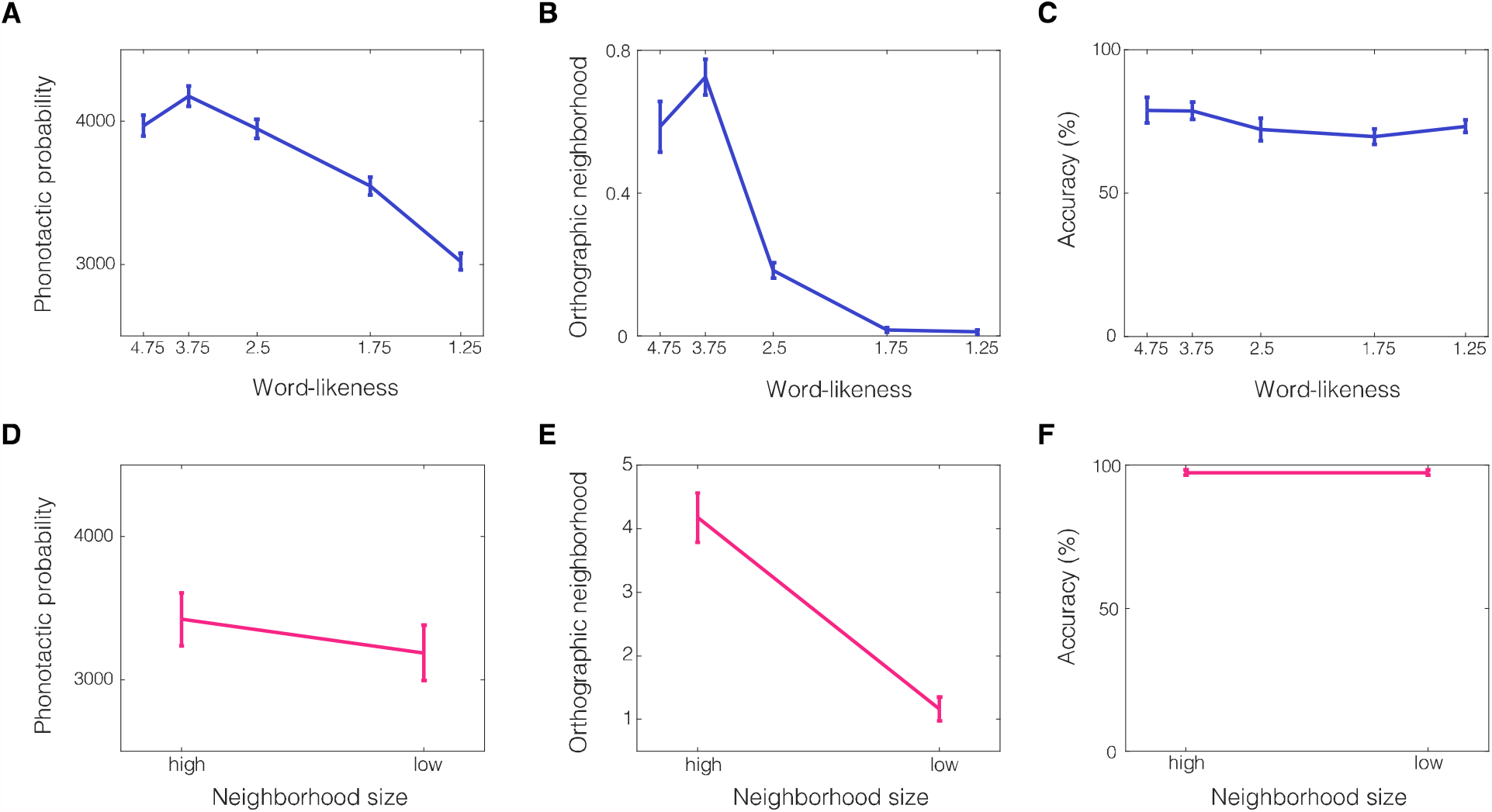
Stimulus characteristics and behavioral results, Experiments 2 (A-C) and 3 (D-F). **A** – Phonotactic probability of the materials in Experiment 2. The y-axis represents phonotactic probability (Methods) - the mean count from an English corpus of all bigrams that occur in a nonword; the x-axis represents the 5 conditions in Experiment 2, ordered by the bin centers of the word-likeless ratings, from most to least word-like. Note that the highest word-like group (bin center 4.75) mostly contained real words, but all 4 other groups contained only nonwords. **B** – Orthographic neighborhood size of the materials in Experiment 2. The y-axis represents orthographic neighborhood size (Methods), i.e., the number of real words that are identical to the nonword up to a substitution of a single letter. The x-axis is the same as in A. **C** - Behavioral results in Experiment 2. The y-axis is the accuracy in the memory probe task (Methods). The x-axis is the same as in A. **D** - Phonotactic probability of the materials in Experiment 3. The y-axis is the same as in A. The x-axis represents the 2 conditions in Experiment 3. The graph shows a numerical decrease of phonotactic probability due to neighborhood size but this effect is not significant (see text). **E** - Orthographic neighborhood size of the materials in Experiment 3. The y-axis is the same as in B. The x-axis is the same as in D. **F** - Behavioral results in Experiment 3. The y-axis is the accuracy in the 1-back task (Methods). The x-axis is the same as in D.

Thus, Experiment 2 suggested that the language system is strongly sensitive to the ‘word-likeness’ of nonwords.

### Key result 3: No evidence for lexical ‘neighbors’ driving the language network’s response to nonwords (Experiments 2 and 3)

One possible explanation for the results of Experiment 2 is that reading nonwords that are word-like activates the representations of real words that are similar to them (e.g., BRIVEARY → BREVIARY or BRAVERY). Given the strong sensitivity of the high-level language network to word meanings (e.g., Fedorenko et al., 2012), stronger responses to more word-like nonwords could be explained on a purely lexical basis, without invoking sub-lexical / phonological regularities.

We tested this possibility in two ways: *first*, in Experiment 3, we measured neural responses to two groups of nonwords that were matched on phonotactic probability (two-sample *t*-test: t(172)=1.1, p=0.27, **Figure 3D**) but differed in the size of their phonological (and orthographic) neighborhood (t(172)=6.9 p<0.001, **Figure 3E**); and *second*, we computed the average orthographic neighborhood size of the nonwords in the five conditions of Experiment 2 and assessed the correlation between this measure and neural response strength.

In Experiment 3, the high- and low-neighborhood conditions elicited activations that were comparable in magnitude across the language network (pink bars in **Figure 2**, Experiment 3), with no evidence for stronger responses to high-neighborhood nonwords across the network (i.e., treating the fROIs as a random effect) or in any of the individual fROIs (*ps*>0.1, **Table 2** and **3**). (Behaviorally, participants were at ceiling on the 1-back task (**Figure 3F**).)

In Experiment 2, of greatest relevance are the two conditions with the lowest word-likeness ratings (the two rightmost, light blue bars in **Figure 2**, Experiment 2). Although these conditions have similarly low orthographic neighborhood size (both around 0; two-sample t-test: t(718)=0.6, p=0.52, **Figure 3B**), they elicited differential brain responses such that the 2nd lowest word-like condition activated the language network significantly more than the least word-like condition (a post-hoc LME revealed a small but significant difference in PSC between these two conditions=0.14, t(158)=2.4, p=0.017). In contrast to their similar orthographic neighborhood size, these conditions differ reliably in their phonotactic probability (t(718)=6.2, p=8e-10; **Figure 3A**), largely mirroring the word-likeness ratings.

In summary, the results of both Experiment 3 and the post-hoc analysis of the two least word-like conditions in Experiment 2 suggest that phonotactic probability (likely reflected in the word-likeness ratings in Experiment 2) explains neural responses in the language regions better than orthographic neighborhood size, and suggest that these responses are not likely to be due to the activation of lexical representations of neighboring real words.

## DISCUSSION

Across five fMRI experiments, we investigated the responses of ‘high-level’ language processing brain regions (e.g., Fedorenko & Thompson-Schill, 2014) to nonwords—meaningless and unconnected strings of sounds/letters (e.g., punes, silory, flope)—and found that these regions indeed respond to such stimuli in an abstract (modality- and task-independent) fashion, suggesting that they represent and process sub-lexical sound-level regularities. In the remainder of the discussion, we situate these findings in the broader theoretical and empirical context and discuss their implications.

### Response of the high-level language network to nonwords

A network of frontal and temporal brain regions supports language processing. These regions respond during both listening to and reading of linguistic stimuli (e.g., Fedorenko et al., 2010; Vagharchakian et al., 2012; Regev et al., 2013; Scott et al., 2017) across tasks (e.g., Fedorenko et al., 2010; Cheung et al., 2020; Diachek et al., 2020), but show little or no response to diverse non-linguistic functions (e.g., Fedorenko et al., 2011; Monti et al., 2012; Fedorenko and Blank, 2020; Ivanova et al., 2020, 2021).

The precise contributions of this network to language processing remain debated (Hickok and Poeppel, 2007; Price, 2010; Friederici, 2011; Indefrey, 2011; Hagoort, 2013; Duffau et al., 2014). Many have argued that distinct subsets of this network store/process syntactic structure vs. word meanings (e.g., Grodzinsky and Santi, 2008; Baggio and Hagoort, 2011; Friederici, 2011, 2012; Tyler et al., 2011; Duffau et al., 2014; Ullman, 2015). However, evidence has been accumulating against this distinction, suggesting that these brain regions support both lexico-semantic and syntactic processing (e.g., Dick et al., 2001; Wilson and Saygin, 2004; Fedorenko et al., 2010, 2012, 2020; Bautista and Wilson, 2016; Blank et al., 2016).

The current study establishes that the language network is sensitive to sub-lexical sound patterns, as evidenced by responses to strings of phonemes that do not constitute real words. The response to nonwords in these brain areas is, by definition, lower than the response to sentences because this network is defined by the *sentences>nonwords* contrast (Fedorenko et al., 2010). Nevertheless, nonwords elicit a response that is consistently and reliably higher than the low-level baseline. Above-baseline responses to nonwords in the language network can be observed in prior fMRI (e.g., Fedorenko et al., 2010; Mahowald and Fedorenko, 2016; Mollica et al., 2020; Chen et al., 2021) and intracranial (e.g., Fedorenko et al., 2016) reports. Additionally, previous data shows that the responses to nonwords are larger than to non-linguistic stimuli such as music and arithmetic tasks (e.g., Fedorenko and Blank, 2020; Chen et al., 2021). However, this is the first study to systematically investigate the responses of the language network to nonwords and try to understand what drives them.

Combined with prior studies, our results suggest that the fronto-temporal language network supports not only the processing of words and inter-word dependencies, but also of phoneme strings that obey phoneme-combinatorial constraints but do not map onto meaningful concepts and are not combinable into phrases. Any proposal about the language network’s computations therefore needs to be able to account for this functional property.

### Modality- and task-independence of the language network’s response to nonwords

The fact that the language regions respond both when participants read nonwords (Experiments 1a, 2, and 3) and when they listen to them (Experiments 1b/c) demonstrates that the representation of nonwords is abstract (unpublished findings from Rebecca Saxe’s lab further show that nonwords in ASL—signs similar in form to meaningful ones but lacking meaning— also elicit above-baseline responses in high-level language areas). These results align with previous findings of modality-independent responses of the language network to stories, sentences, and word lists (e.g., Fedorenko et al., 2010; Vagharchakian et al., 2012; Regev et al., 2013), but critically extend them to stimuli that lack meaning.

Similarly, our results demonstrate that the language regions respond to nonwords across tasks, in line with the task-independence of the language network’s responses to words and sentences (e.g., Cheung et al., 2020; Diachek et al., 2020). Past neuroimaging and patient studies investigating phonological processing have used artificial tasks such as rhyme judgements (e.g., Petersen et al., 1989; Paulesu et al., 1993; Seghier et al., 2004; Geva et al., 2011; Pillay et al., 2014; Yen et al., 2019), nonword repetition (e.g., Fridriksson et al., 2010; Church et al., 2011; Scott and Perrachione, 2019), active maintenance of words/nonwords in memory (e.g., Paulesu et al., 1993; Awh et al., 1996), or phoneme discrimination/identification (e.g., Démonet et al., 1992; Zatorre et al., 1992; Burton et al., 2000). Such tasks differ substantially from the natural ‘task’ we perform when faced with linguistic input—to extract meaning. Artificial-task-based paradigms may engage cognitive processes beyond those that support the processing of linguistic input and are instead related to the task demands, including executive processes supported by the domain-general multiple demand network (e.g., Diachek et al., 2020; Fedorenko and Shain *in prep*) or the motor articulation system (e.g., Bohland and Guenther, 2006; Basilakos et al., 2017).

In contrast, we show that even passive listening to or reading of nonwords elicits a response in the high-level language network, suggesting that the responses we observe reflect something about the *intrinsic computations* necessary for recognizing and processing sub-lexical sound patterns. Such computations are presumably critical to language processing and acquisition given that any new word we encounter is at first just a sequence of sounds that gradually acquires semantic associations as we learn the word’s meaning (e.g., Davis et al., 2009; Perry et al., 2018; Jones et al., 2021).

### Sensitivity of the language network to phonotactic probability of nonwords

In Experiment 2 we found that more word-like/phonotactically probable nonwords elicit stronger responses in the language network. This result plausibly reflects a process of matching sounds/sound patterns to stored representations extracted from our lifetime of experience with a language, whereby the strength of response is proportional to how well the stimulus matches stored linguistic regularities and the amount of matching information (e.g., Hayes and Wilson, 2008). This representation is consistent with the notion of ‘phonological schemata’ (Jackendoff, 2002). Storage of frequent sound/letter n-grams allows for more efficient processing through enabling the representation assembly to proceed in larger chunks than single phonemes/letters (Bybee, 1999; Bybee and Hopper, 2001; Vitevitch and Luce, 2005; O’Donnell, 2015).

Vinckier et al. (2007) reported stronger responses to more frequent letter combinations in the occipito-temporal orthographic pathway (which their study focused on) with a posterior-to-anterior progression, and this contrast became even stronger in frontal and temporal brain areas that plausibly correspond to the high-level language areas investigated here. However, those areas were not functionally defined in their study, making it impossible to unambiguously interpret the results (Poldrack, 2006; Fedorenko, 2021).

Might stronger responses to more word-like nonwords reflect activation of lexical representations of real words that share phonological/sound structure with them? We evaluated this possibility in two ways, and did not find support for it. In Experiment 3, nonwords that differed in the number of real-word neighbors (but were matched on phonotactic probability) elicited similar-magnitude responses in the language network. And in Experiment 2, we found that although the two least word-like groups of nonwords both had few/no real-word neighbors, the more word-like nonwords elicited stronger responses in the language network. So, it appears that the language regions represent sub-lexical units including phoneme sequences that do not have a lexical-semantic representation, and that the frequency of these phoneme sequences in our experience with the language is what drives the response to nonwords in the language regions, even if familiar sound patterns do not lead to activation of similar-sounding real words.

However, a possibility that is more difficult to rule out is that the response to nonwords is, at least in part, driven by semantic associations that might be elicited by particular sounds/sound clusters (e.g., Iwasaki et al., 2007; Monaghan et al., 2014; Larsson, 2015; Blasi et al., 2016; Winter et al., 2017; Sidhu and Pexman, 2018; Pimentel et al., 2019; Vinson et al., 2021) or morphemes/morpheme-like elements that occur in some nonwords. Further research is needed to determine whether sub-lexical semantic associations may be sufficient to explain the response to nonwords.

## Conclusions and limitations

We have presented evidence that meaningless sequences of phonemes—auditory or visual— elicit responses in the network of brain regions that has been traditionally associated with the processing of word meanings and word-combinatorial processing. This finding aligns with views of linguistic knowledge and processing where no sharp boundaries are drawn between phonemes, morphemes, words, and higher-order units like constructions (e.g., Gaskell and Marslen-Wilson, 1997; Bybee, 1999, 2013; Jackendoff, 2007; Huettig et al., 2020; Jackendoff and Audring, 2020), and challenges accounts of the language network, or its subcomponents, that focus on compositional meaning, or prediction at the level of word sequences.

More research is needed to understand the precise features that make a nonword elicit an above-baseline response, including potential contributions from semantic associations elicited by sub-lexical units, as discussed above. In addition, developmental investigations, especially during the first few years of life—when most words we encounter do not yet have meaning—could help illuminate the formation of linguistic knowledge representations (e.g., Jones et al., 2021).

## Acknowledgements

We would like to acknowledge the Athinoula A. Martinos Imaging Center at the McGovern Institute for Brain Research at MIT, and its support team (Steve Shannon and Atsushi Takahashi). We thank former and current EvLab members for their help with fMRI data collection, Steve Piantadosi for help in creating the materials for Experiment 2, Peter Graff for early discussions of phonology in the brain, Alex Paunov for help with some of the analyses, and Ray Jackendoff for helpful comments on this work. TIR also thanks Janet Werker, the audience at the 2020 Neurobiology of Language conference, and Tali Bitan and her lab for helpful discussions. TIR was supported by the Zuckerman-CHE STEM Leadership Program. EF was supported by the R00 award HD057522, R01 awards DC016607 and DC016950, and funds from the Brain and Cognitive Sciences department and the McGovern Institute for Brain Research.

